# Enterobacteriaceae isolated from patients share antibiotic resistance conferring plasmids recently acquired from those isolated from sinks in the same treatment room

**DOI:** 10.1101/2022.11.01.514792

**Authors:** Bradford P Taylor, You Che, Hemanoel Passarelli, Gill Smollan, Carmit Cohen, Rotem Rapaport, Ilana Tal, Nani Pinas Zade, Hanaa Jaber, Nati Keller, William P Hanage, Gili Regev-Yochay

**Author notes:** These authors contributed equally (William P Hanage, Gili Regev-Yochay).

## Abstract

Identifying how and where pathogens acquire antibiotic resistance is crucial to developing effective strategies to limit its spread. Many bacterial species carry and share plasmids harboring antibiotic resistant genes. Plasmids are mobile genetic elements whose horizontal transmission is difficult to assess through genomic comparison due to assembly issues when using short-read sequencing alone. In this study, we use hybrid assembly to fully assemble plasmids that are shared between different Enterobacteriaceae isolated from patients and sinks in the same hospital rooms. We isolated and sequenced pairs of carbapenem resistant *Enterobacter hormaechei* subsp. *xiangfangensis* and *Klebsiella pneumoniae* from patients and sinks within the same hospital room. The isolate pairs share plasmids that putatively confer antibiotic resistance, including carbapenem resistance. These plasmids differ by few mutations and structural changes, while the isolates carry unique plasmids. Together, this suggests that plasmids can act as vectors of antibiotic resistance spread from sink reservoirs to patients.

## Introduction

Antibiotic resistance poses a growing threat to modern medicine by reducing the effective treatment options against bacterial infections (World Health Organization 2014). This threat is particularly acute in hospitals where infections by resistant pathogens arise from nosocomial spread as they can persist across different hospital environmental reservoirs seemingly for months at a time (Carling 2018, Constantinides 2020, David 2019). Given how widespread resistant pathogens are within hospitals, it is essential to identify how antibiotic resistance spreads.

Comparing sequences derived from bacterial isolates, such as for antibiotic resistance genes, allows one to identify differences which can be used to estimate the recency of divergence between the two populations. For Gram-negative bacteria, resistance genes mainly reside on plasmids – extrachromosomal DNA capable of both horizontal transfer via conjugation and vertical transmission via reproduction (Carattoli 2013). Independent horizontal transmission of resistant loci means that establishing transmission can only be confidently done by direct comparison of plasmids.

Direct comparison of full plasmid sequences is only recently feasible thanks to advances in sequencing and assembly methods. Short-read sequencing provides high coverage and less perbase error, but it often fails to assemble complete plasmids because ambiguities may arise when assembling long stretches of repeat regions common in plasmids (Klassen 2012). In contrast, long-read sequencing generate reads that span across these repeat regions, but it is usually limited by the low coverage and high per-base error rate (Lu 2016). Therefore, combining short- and long-read sequencing of the same sample allows hybrid assembly that overcomes the deficiencies of each individual method.

Here, we show recent divergence between plasmids shared between *Enterobacteriaceae* isolated from patients and sinks in their same treatment rooms. We detail the resistance profiles of the shared plasmids, which includes carbapenem resistance genes. We analyze the genetic divergence between shared plasmids and show that plasmid acquisition is recent, relative to an estimated mutation rate. Overall, this work shows how full plasmid sequences obtained through hybrid assembly can help resolve divergence rates and, for specific pairs of plasmids, nosocomial transmission of resistance genes.

## Results

We sampled and sequenced individual colonies from four carbapenem-resistant positive cultures isolated from concurrent pairs of patients and sinks within the same hospital room. Hybrid assembly resolved the chromosome and 3 plasmids within each sample except for one sample where the genome was split into 15 contigs including 2 fully circular plasmids. One room contained *Klebsiella pneumoniae* ST11 isolates and another contained *Enterobacter hormaechei* subsp. xiangfangensis ST 136. We circularized the split *E. hormaechei* chromosome in the sink sample using RagTag (Alonge et al 2021). Overall, the paired isolates share similar sequences.

Figure 1 shows local alignment of contigs between isolates from the same hospital room. The ribbon plots show that room pairs share chromosomes and plasmids with high sequence identity and synteny, but that there are also plasmids unique to each sample. The darker ribbons connect regions that statistically share sequence identity, but may not necessarily reflect structural variation. Dot plots generated using mummer corroborate the lack of structural variation with only one inversion observed between the 157 kb plasmid shared between *Enterobacter* isolates (Supplementary Figure 1). We summarize global features of the contigs initially output by unicycler in Table 1 and color code the putatively shared plasmid pairs between isolates.

**Figure 1:**
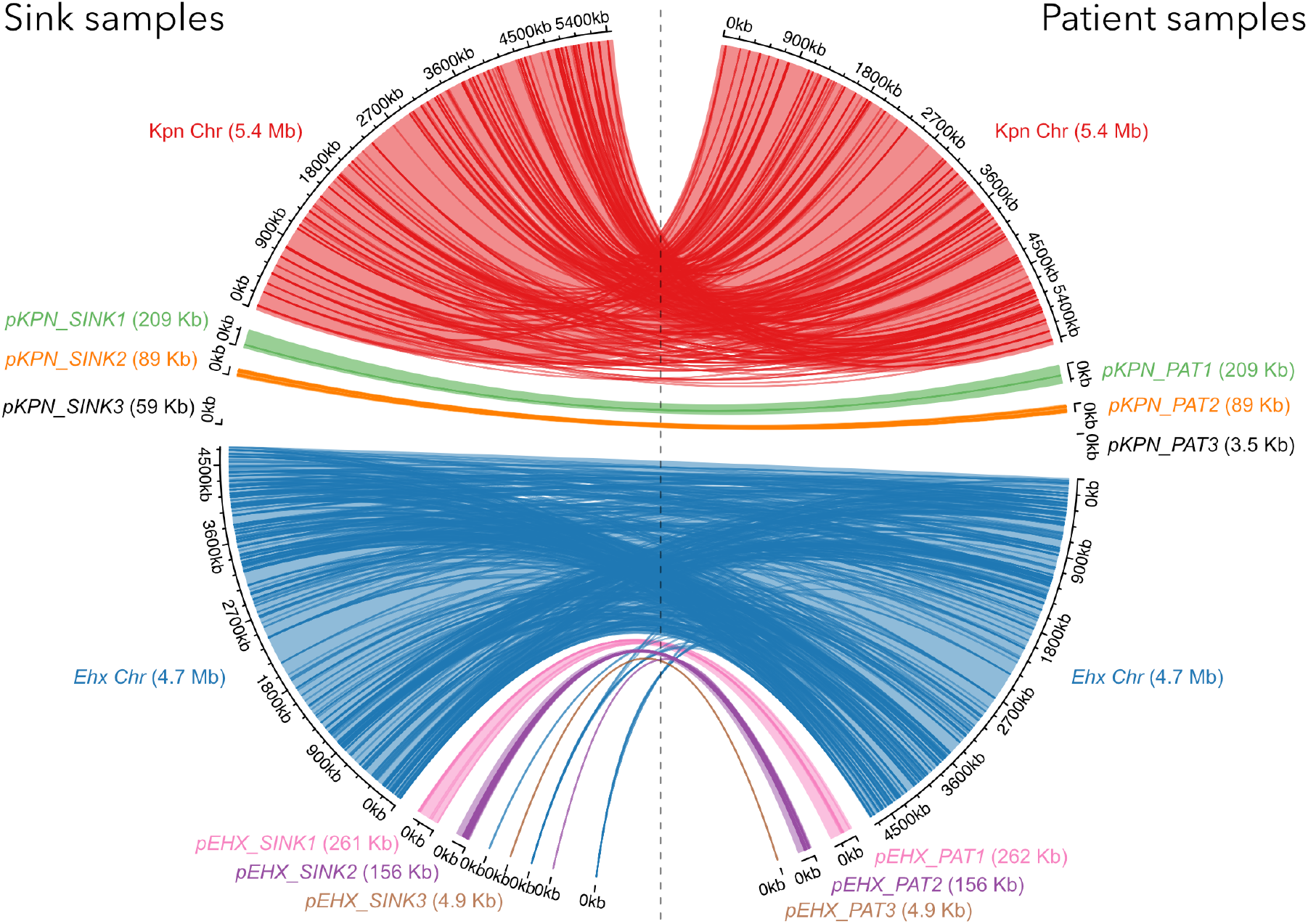
Locally aligned contigs from four Enterobacteriaceae isolates. Ribbons connect locally aligned regions. We performed local alignments only between isolates obtained from the same hospital room.

| Sample | Contig | Length | GC content | CDS | Coding length | Coding density | Inc type | depth | circular |
| --- | --- | --- | --- | --- | --- | --- | --- | --- | --- |
| Klebsiella Sink | 1 | 5456289 | 0.574 | - | - | - |  | 1.00x | O |
|  | 2 | 209198 | 0.452 | 226 | 166271 | 0.7948020536 | IncHI | 1.68x | O |
|  | 3 | 89653 | 0.523 | 106 | 74458 | 0.8305132009 |  | 2.12x | O |
|  | 4 | 59140 | 0.521 | 79 | 58673 | 0.9921034833 |  | 3.12x | O |
| Klebsiella Patient | 1 | 5456077 | 0.574 | - | - | - |  | 1.00x | O |
|  | 2 | 209215 | 0.452 | 223 | 166170 | 0.7942547141 | IncHI | 1.79x | O |
|  | 3 | 89653 | 0.523 | 107 | 74586 | 0.8319409278 | IncA | 2.16x | O |
|  | 4 | 3560 | 0.513 | 3 | 1851 | 0.5199438202 |  | 16.51x | O |
| Enterobacter Sink | 1 | 3293535 | 0.552 | - | - | - |  | 1.00x |  |
|  | 2 | 1006754 | 0.554 | - | - | - |  | 1.27x |  |
|  | 3 | 410285 | 0.561 | - | - | - |  | 1.08x |  |
|  | 4 | 261708 | 0.461 | 285 | 222704 | 0.8509636694 | IncHI | 1.31x |  |
|  | 5 | 156916 | 0.534 | 174 | 135902 | 0.8660812154 | IncA | 1.61x | O |
|  | 6 | 6463 | 0.569 | 8 | 5986 | 0.9261952654 |  | 1.82x |  |
|  | 7 | 4995 | 0.518 | 8 | 2665 | 0.5335335335 |  | 13.94x | O |
|  | 8 | 1635 | 0.534 | 0 |  | 0 |  | 1.27x |  |
|  | 9 | 1270 | 0.494 | 2 | 1011 | 0.7960629921 |  | 1.31x |  |
|  | 10 | 920 | 0.539 | 0 |  | 0 |  | 1.19x |  |
|  | 11 | 343 | 0.484 | 0 |  | 0 |  | 0.76x |  |
|  | 12 | 154 | 0.403 | 0 |  | 0 |  | 0.51x |  |
|  | 13 | 147 | 0.456 | 0 |  | 0 |  | 1.27x |  |
|  | 14 | 110 | 0.536 | 0 |  | 0 |  | 0.51x |  |
|  | 15 | 106 | 0.34 | 0 |  | 0 |  | 1.47x |  |
| Enterobacter Patient | 1 | 4726467 | 0.554 | - | - | - |  | 1.00x | O |
|  | 2 | 262065 | 0.461 | 285 | 222717 | 0.8498540438 | IncHI | 1.33x | O |
|  | 3 | 156088 | 0.534 | 173 | 135339 | 0.8670685767 | IncA | 1.89x | O |
|  | 4 | 4995 | 0.518 | 9 | 2889 | 0.5783783784 |  | 6.16x | O |

We annotated the contigs using PGAP to identify potential phenotypes carried on the plasmids which may be transferred between sink and patient isolates. Figure 2 visualizes the shared plasmids highlighting coding sequences (CDS) that putatively confer antibiotic resistance and plasmid mobility. These plots focus on the patient plasmids, but are representative of the paired sink plasmids. Overall, there are no major structural changes between pairs except a single inversion which we highlight on pEHX_PAT2.

**Figure 2.**
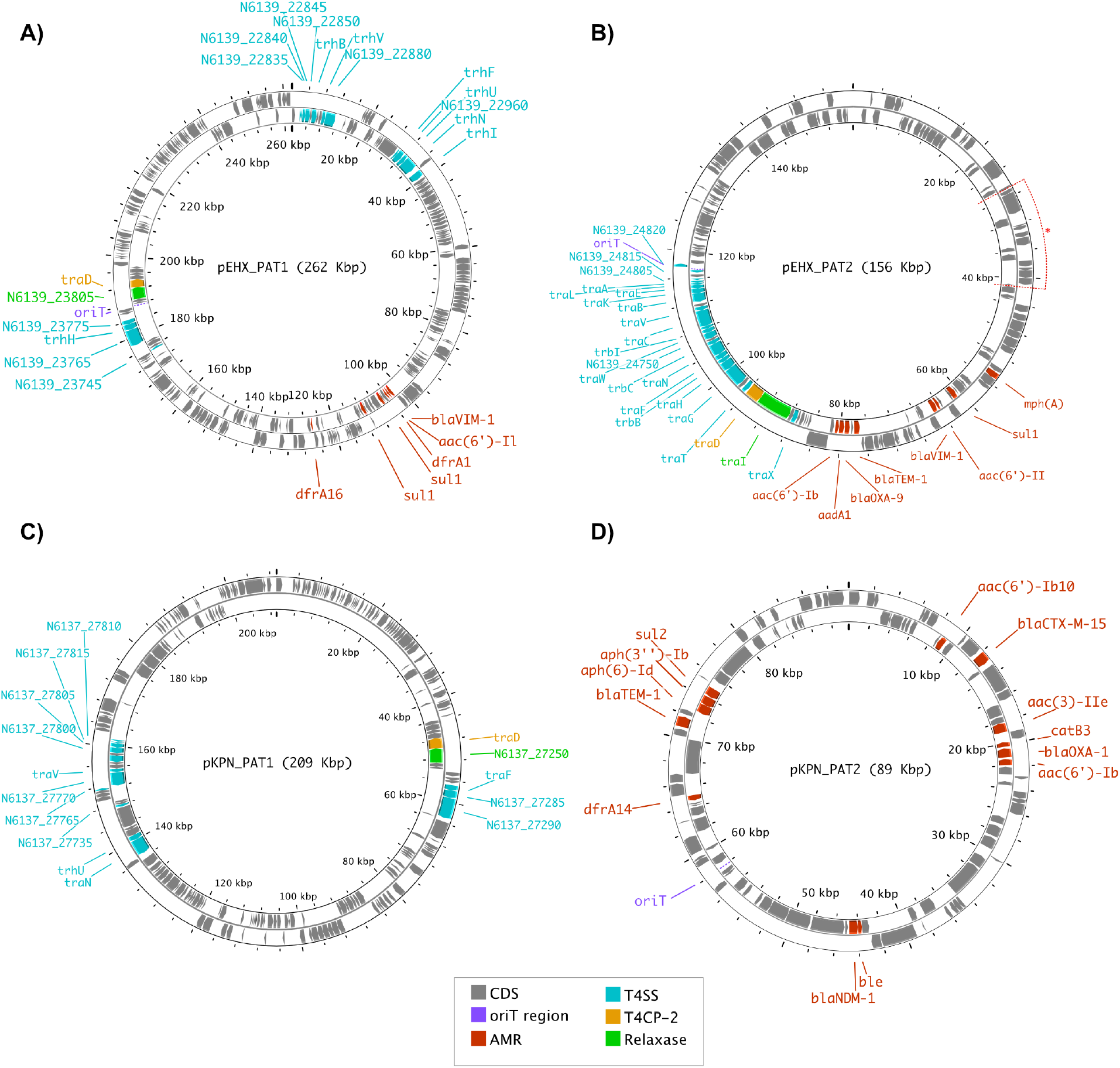
Plasmids shared between sink and patient isolates. We highlight genes associated with antibiotic resistance (red) and plasmid mobility (all other non-grey colors). The outer ring contains CDS on the positive strand and the inner ring contains CDS on the negative strand. The inversion between the pEHX_PAT2/pEHX_SINK2 pair is highlighted with red wedge marked “*”.

All shared plasmids, except the larger *K. pneumoniae* plasmid pKPN_PAT1/pKPN_SINK1, putatively confer antibiotic resistance as identified using AMRFinder. The smaller shared Klebsiella plasmid, pKPN_PAT2/pKPN_SINK2, putatively contains the broadest resistance profile including aminoglycosides (*aac(6′)-Ib, aac(3)-IIe, aac(6′)-Ib-cr5, aph(6)-Id, aph(3″)-Ib)*, beta-lactams (*bla_CTC-M-15_, bla_OXA-1_, bla_NDM-1_, bla_TEM-1_*), bleomycin (*ble*), trimethoprim (*dfrA14*), and sulfonamides (*sul2*). There are a large number of transposases on this plasmid which likely contribute to the formation of such a broad resistance profile. This plasmid is mobilizable (*MobC),* but lacks conjugative machinery, and likely relies on another plasmid for transmission.

Both plasmid pairs shared between *E. hormaechei* isolates putatively confer antibiotic resistance. These pairs both putatively confer beta-lactam (*bla_VIM-1_*), aminoglycoside (*aac(6′)-Il*) and sulfonamide (*sul1*) resistance from the same *loci*. In particular, *bla_VIM-1_* and *aac(6′)-II* are adjacent to each other and to transposases on each plasmid, which may reflect horizontal gene transfer of resistant loci between them. Each pair of shared Enterobacter plasmids carry resistance loci unique to the pair. The larger shared plasmid, pEHX_PAT1/pEHX_SINK1 uniquely, putatively carries resistance to trimethoprim (*dfrA1, dfrA16*) and quinolones (*qnrA1*). The smaller shared plasmid pEHX_PAT2/pEHX_SINK2 uniquely, putatively carries resistance to aminoglycosides (*aac(6′)-Ib′, aadA1*), beta-lactams (*bla_TEM-1_, bla_OXA-9_*), and macrolides (*mph(A)*). The genomic inversion between the smaller Enterobacter plasmid pair does not include any resistance loci.

Lastly, we aimed to assess the total genetic divergence between shared plasmids through local alignment. For the smaller Enterobacter plasmid pair, we separated and reverse-complemented the inversion and individually aligned the resulting three portions of the plasmids. In that case, we further ignored the ends with long stretches of gaps that result from difficulty in alignment given structural variation, including a difference in copy number of an IS-26 transposase. Each *K. pneumoniae* plasmid differ by 2 SNPs with its respective pair giving a SNP frequency of 1.3E-5 for the pKPN_PAT1/pKPN_SINK1 and 2.2E-5 for pKPN_PAT2/pKPN_SINK2. The larger Enterobacter pair, pEHX_PAT1/ pEHX_SINK1, differ by 8 SNPs; however, 7 SNPs occur within a 500-bp neighborhood which may reflect a recombination event. The smaller *Enterobacter* pair, pEHX_PAT2/ pEHX_SINK2, contains 3 SNPs with 2 occurring within 5-bp. Nearby SNPs may indicate recombination events as opposed to independent SNPs. Treating SNPs that occur within 500-bp of another SNP as a single mutational event gives mutation frequencies of 7.6E-6 for pEHX_PAT1/ pEHX_SINK1 and 6.4E-6 for pEHX_PAT2/ pEHX_SINK2. We caution against overinterpreting these differences, as they are calculated from a small number of SNPs. In practice, it is impossible to reliably compare rates in order to distinguish whether shared plasmids were acquired during a single conjugation event or sequentially during multiple conjugation events. However, this indistinguishability suggests the pairs of plasmids were shared recently as compared to the rate of SNP accumulation.

## Discussion

Here, we used hybrid assembly to genetically compare plasmids recently shared between pairs of *Enterobacteriaceae* isolated from patients and sinks in the same room. Structural variant and mutational analysis show that the shared plasmids diverged by only a small number of mutations and recombination events. Our work suggests the sink acted as a reservoir for recent nosocomial transmission of antibiotic resistance via plasmids, given patients only transiently occupy the room.

While we limited our study to just a few isolates, this work demonstrates the power of hybrid assembly to fully resolve the plasmid sequences to allow comparison to assess transmission routes and rates. While prior studies of nosocomial antibiotic resistance spread were larger in size, they focused mostly on identifying the presence of resistance either using culturing or shortread sequencing (although see (Chen 2021) where they identified a <10kb plasmid shared between samples). Further sampling can lead to broader analysis that potentially resolve networks of dispersal within the hospital. By understanding the plasmid biogeography within hospitals, we can inform infrastructure and infection control practices aimed at reducing the spread of resistance to patients.

## Methods

### Isolate collection

We preferentially selected 4 carbapenem resistant isolates within a broader surveillance at Sheba Medical Center in central Israel. Sink screening was performed by swabbing the sink-outlet surface with 4 sterile cotton swabs or inserting the 4 swabs as far as possible into the sink-outlets and similarly swabbing the surface of the pipe leading to the sink-trap. To avoid enrichment of Enterobacteriaceae on expanse of other bacterial species, swabs were placed in sterile saline and transferred immediately to the laboratory. Samples were isolated on 2018-10-13 for the *Klebsiella* pair and 2018-01-01 for the *Enterobacter* pair. Patient samples were obtained from rectal swabs using Copan Amies sterile transport swabs (Copan Diagnostics). Swabs were streaked in the classical method onto Chromagar KPC plates (Hy Laboratories, Rehovot) to achieve isolated colonies and incubated overnight at 35°C in ambient air. The containers were vortexed vigorously prior to streaking the liquid onto Chromagar KPC plates with a new sterile swab. Suspicious colonies were identified using Maldi-TOF. Carbapenemase genes were identified by PCR using Xpert Carba-R cartridges (GeneXpert Cepheid).

### Whole genome sequencing and annotation

The Oxford Nanopore library preparation was performed using the Oxford Nanopore Technologies 1D ligation sequencing kit SQK LSK109 (Oxford Nanopore, Oxford, U.K.), according to the manufacturer’s instructions. Sequencing was performed on the MinION MK1 using a R9.4 flow cell. The Illumina libraries were prepared using Nextera XT (Illunima, San Diego, CA, USA). Samples were sequenced using the Illumina Miseq. We performed hybrid assembly using the default settings of unicycler v0.4.8. We circularized the split *E. hormaechei* chromosome using RagTag with Enterobacter hormaechei subsp. xiangfangensis LMG27195 (type strain) (NZ_CP017183.1) and Enterobacter hormaechei subsp. xiangfangensis 34399 (NZ_CP010384.1) as references (Alonge et al 2021). Sequences were uploaded to NCBI and assigned the accession numbers: CP106898-CP106905, CP106894-CP106897, CP106890-CP106893, and CP106886-CP106889.

We classified isolates by comparing chromosomal contigs to the JSpecies database (Richter et al 2016). We annotated all circular plasmid contigs using PGAP (Li et al 2021). Sequence types (STs) were assigned employing MLST v2.0.9 using the *K. pneumoniae* MLST scheme and *Enterobacter cloaceae* schemes (Larsen et al 2012). We identified antibiotic resistance *loci* using AMRfinder. We identified plasmid mobility loci using oriTfinder (Li et al 2018).

### Alignment

We locally aligned paired scaffolds using BLAST v2.5.0 (Altschul et al 1990). For SNP comparison, shared plasmids were aligned to each other first using nucmer V4.0.0rc1 to assess structural variation (Marçais et al 2018). SNPs were identified by aligning shared plasmids using mafft after manually rearranging any inversions (Katoh et al 2013). The plasmid pair with large structural differences, contiguous regions of indels and inversions, were separated into non-inversion and inversion regions before aligning with mafft. Only SNPs that occurred in regions absent of long stretches of gaps were considered during analysis in order to ignore alignment issues arising due to the indels.

## Supplementary Information

**Supplementary Figure 1:**
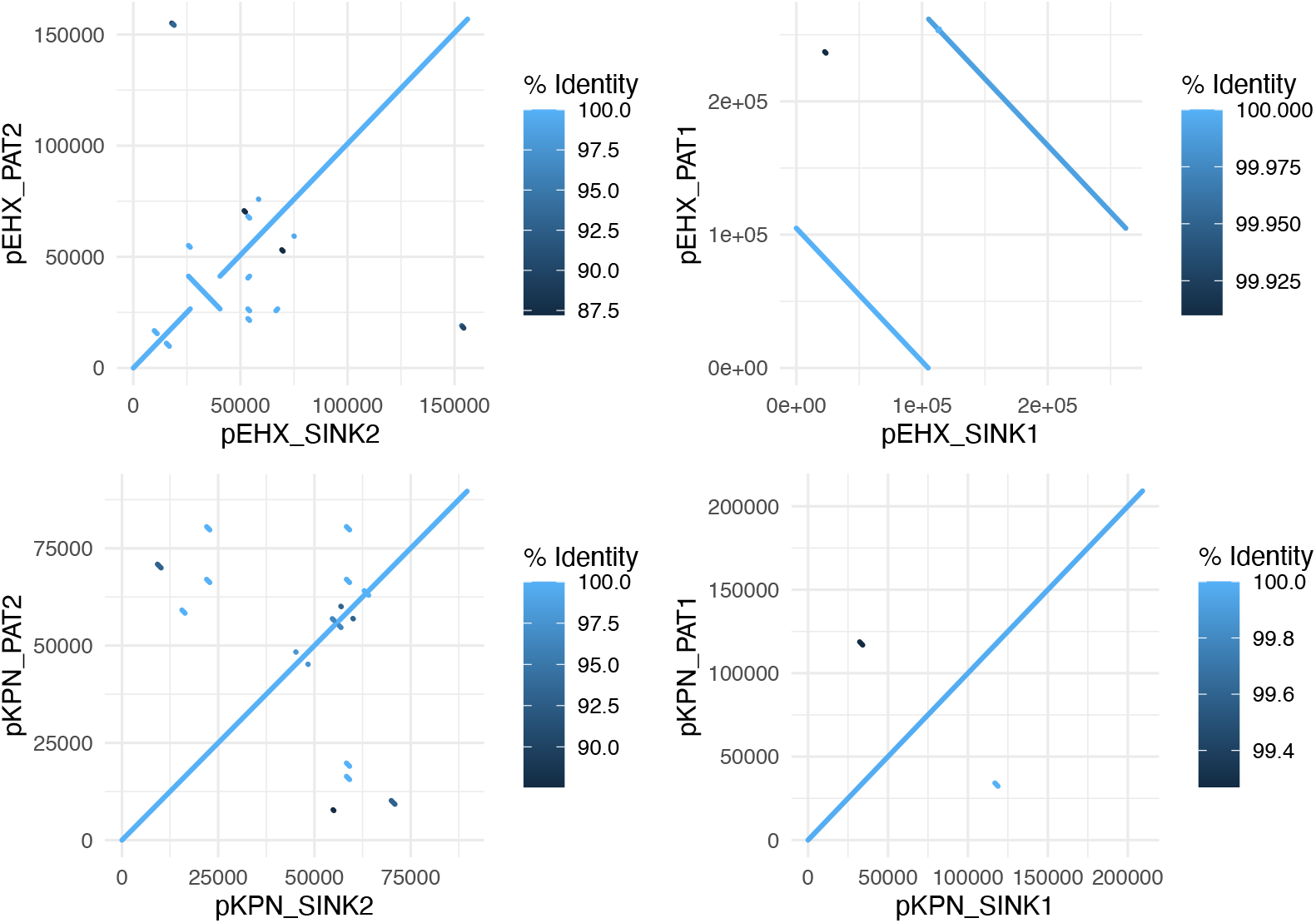
Dot-plot comparison of similarly sized plasmid contigs globally aligned using Mummer. Diagonal lines indicate regions of alignment using a 65-bp resolution. Lines perpendicular to the diagonal show alignment between regions that are reverse complements between sequences--a hallmark of inversions. Off-diagonal lines can indicate transposition of regions across the genome, but may also arise spuriously due to sequence similarity and reverse complementary similarity.

